# A novel haplotype-based eQTL approach identifies genetic associations not detected through conventional SNP-based methods

**DOI:** 10.1101/2020.07.23.206391

**Authors:** Ziad Al Bkhetan, Gursharan Chana, Cheng Soon Ong, Benjamin Goudey, Kotagiri Ramamohanarao

## Abstract

**Motivation:** The high accuracy of current haplotype phasing tools has enabled the interrogation of haplotype (or phase) information more widely in genetic investigations. Including such information in eQTL analysis complements SNP-based approaches as it has the potential to detect associations that may otherwise be missed.

**Results:** We have developed a haplotype-based eQTL approach called *eQTLHap* to investigate associations between gene expression and haplotype blocks. Using simulations, we demonstrate that eQTLHap significantly outperforms typical SNP-based eQTL methods when the causal genetic architecture involves multiple SNPs. We show that phasing errors slightly impact the sensitivity of the proposed method (< 4%). Finally, the application of eQTLHap to real GEUVADIS and GTEx datasets finds 22 associations that replicated in larger studies or other tissues and could not be detected using a single-SNP approach.

**Availability:** https://github.com/ziadbkh/eQTLHap.

## 1 Introduction

Genome-wide association studies (GWAS) have revealed numerous significant genetic associations with diseases. A large number of these variations can not be directly linked to a particular gene [1] as they are not part of the protein-coding regions [2]. Therefore, understanding how they influence a specific disease is a challenging task [3]. One mechanism underlying such associations is that genetic variations, especially those within regulatory regions, can affect gene transcription levels, reducing their functionality or deactivating them completely [4, 3]. As an example, around 50% of GWAS variations associated with schizophrenia have an impact on the expression of related genes [4]. Such cases are investigated statistically through expression quantitative trait loci (eQTL) analysis [5]. Numerous eGenes (genes whose regulation is influenced by SNPs) have been revealed through eQTL analysis applied to multiple populations and tissues [6, 3].

eQTL analysis has been widely applied using a range of approaches, primarily examining the relationship of single SNPs and gene expression [7]. In addition, joint analysis of multiple SNPs, interactions between SNP (epistasis) and phased haplotypes have also been considered [8, 9, 10]. Of these, haplotype-based approaches are amongst the least explored. However, it is well known that expression can be influenced by the phase of the mutations and the gene of interest. Such scenarios include compound heterozygosity, where a disorder is associated with two alternate alleles allocated on different homologous copies of a specific region, as well as allele-specific expression, where the allocation of mutations on each haplotype copy can have a different impact on the expression of homologous gene copies. Hence, phase-aware eQTL analysis is likely to have greater power than SNP approaches with respect to such cases [11].

A barrier for phased haplotype-based eQTL analysis is the increased complexity of the analysis. Many large eQTL studies rely on SNP array data and hence phase need to be estimated, a process which may introduce errors into the analysis. Moreover, decisions also need to be made about how to form haplotype blocks and how to represent these blocks when evaluating their associations with gene expression levels. We have shown that when dealing with haplotype blocks, the choice of partitioning method has an impact on the phasing errors [12]. Previous approaches using haplotypes for eQTL analysis have defined blocks from pairs of SNPs [9], combinations of up to 4 SNPs [10] or regulatory regions [13] but it is unclear how the accuracy of the determined haplotypes is influenced by these partitioning approaches. In contrast, recent work has shown that existing phasing tools can achieve high accuracy within the haplotype blocks defined using linkage disequilibrium (LD) [14, 12]. Such results encourage using LD-based blocks in this context. In addition, the use of LD will also minimise the diversity of the haplotypes within each block, leading to an increased statistical power to uncover associations with phenotypes of interest.

In this study, we present a method for haplotype-based eQTL analysis, called *eQTLHap*. Individuals’ phased haplotypes are partitioned into variable-length blocks based on the LD between SNPs. Using simulated genotype and gene expression data, we compare the detected significant associations when considering individuals’ haplotypes to the ones detected using standard eQTL analysis (single SNP-based) and also considering the block genotype (combining all block’s SNPs). The latter comparison demonstrates the importance of including the allele allocation in the analysis as both approaches consider the same combination of variants. Furthermore, we report the impact of phasing errors on eQTL results for different switch error rates (from 0% to 2.5%). Such novel analysis is essential to demonstrate the reliability and credibility of obtained results in real applications using available computationally phased haplotypes data. Finally, we applied our approach to two real genotype and gene expression datasets, GEUVADIS [3] and GTEx [6] and we investigate the revealed associations.

The results demonstrate the efficacy of the proposed approach for particular genetic architectures underlying the variation of gene expression. Considering phase information in eQTL analysis increases the true positive rate (TPR) of the detected eGenes, primarily when the causal genetic architecture involves multiple SNPs. The three eQTL approaches agree on a large percentage of the associations, yet each captures its own unique subset. There is a slight impact of phasing errors (< 2.5%) on the TPR obtained by our method. eQTLHap uncovers associations (replicated in GTEx and GEUVADIS datasets) in blood that could not be detected by single SNPs but have been replicated in recent meta-analyses. These results highlight the value haplotype-based approached to complement current genotype-based methods for uncovering novel eQTLs.

## 2 Methods

### 2.1 Block determination and encoding

eQTLHap investigates haplotype blocks to conduct eQTL analysis. The blocks are encoded using three different ways depending on the conducted assessment SNP, block’s genotype (denoted as B-Gen) or block’s haplotype (denoted as B-Hap) as illustrated in Figure 1. SNPs are encoded considering an additive impact as the dosage of minor/reference allele as illustrated in Figure 1 b). This SNP encoding is similar to the standard SNP representation in most genetic problems and it does not account for the combined impact of multiple SNPs. The genotypic representation of a block is encoded by concatenating the genotypes of all SNPs within the block, and it is considered as a categorical variable as illustrated in Figure 1 c). This encoding accounts for the combined impact of multiple SNPs. Finally, the haplotypic representation of a block is represented using a bag-of-haplotypes encoding similar to the bag-of-words (BOW) encoding used in text mining [15]. The block of each individual is encoded as the dosage of each possible haplotype across all individuals as illustrated in Figure 1 d). This encoding accounts for both the combined impact of multiple SNPs as well as the allele location that is ignored by the genotypic encoding.

**Figure 1:**
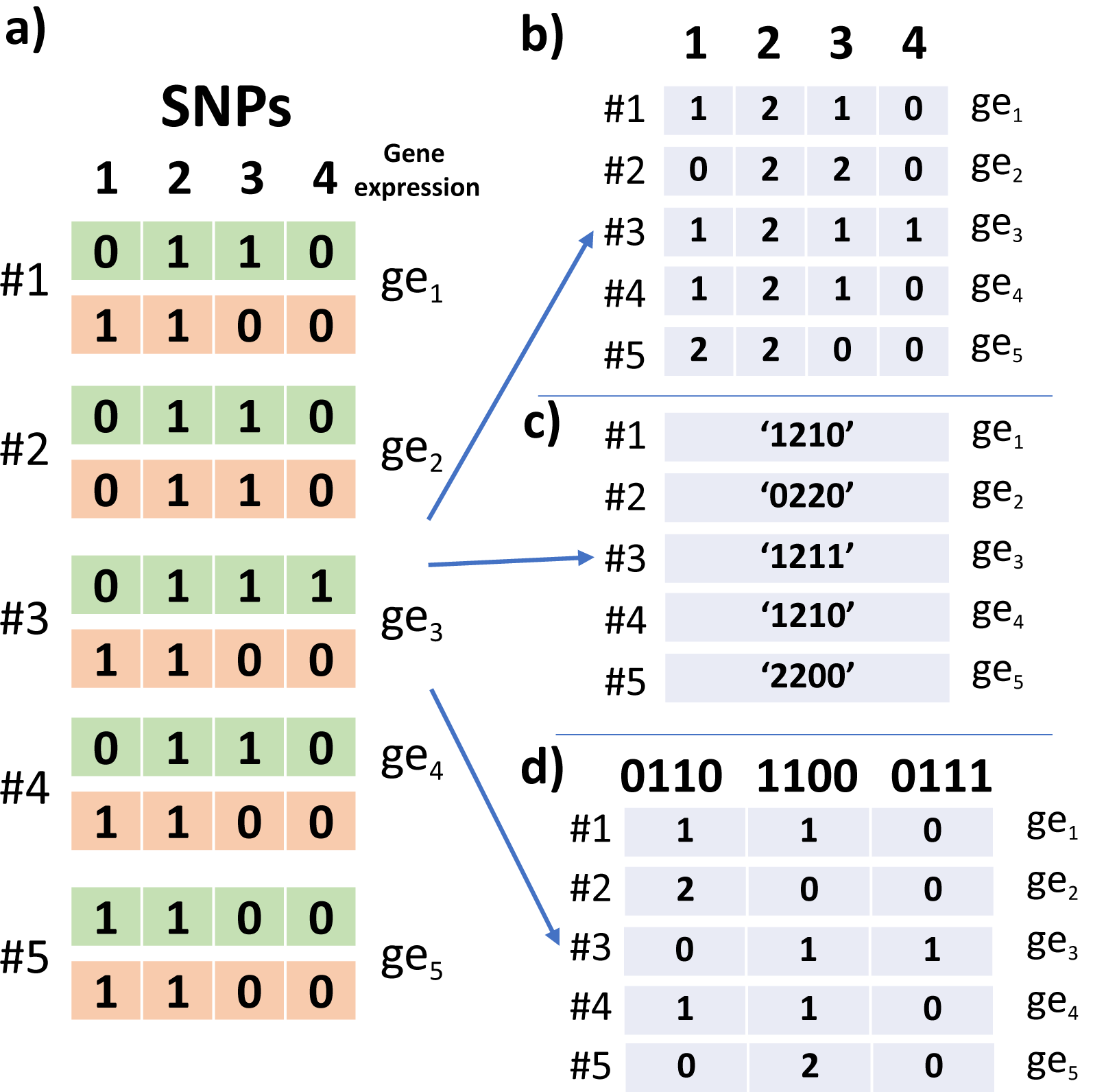
Different encoding for blocks. a) An example representing a block of 4 phased SNPs across 5 individuals and gene expression. Haplotypes are encoded using the dosage of the minor allele within the SNP (0: reference, 1: minor). b) SNPs are represented using their genotypes. c) B-Gen: Block’s genotype is encoded by concatenating the genotypes of block’s SNPs. d) B-Hap: Bag-of-haplotypes encoding for this block as the dosage of the three unique haplotypes within the block.

Haplotype blocks can be determined by several methods such as D-prime confidence interval [16], Four gamete [17], Solid spine [18], big-LD [19] and a simple sliding window. In this study, we used haplotype blocks determined based on LD though PLINK software [20] that implements the Confidence Interval (CI) algorithm [16] (parameters: *–blocks no-pheno-req*). However, our eQTLHap accepts any kind of blocks (overlapping/non-overlapping, fixed or dynamic lengths) as long as the start and end SNP are determined for each block.

### 2.2 Association analysis

eQTLHap relies on applying eQTL analysis based on haplotype blocks and considering the alleles of both chromosome copies. After block determination and encoding as mentioned above, a multiple linear regression model is fitted for gene expression and the haplotype encoding, then *R*^2^, F-test and p-value are calculated:

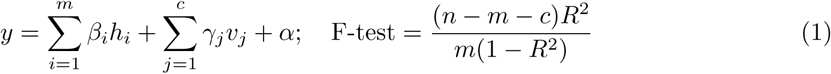

where *y* is the gene expression. *m* is the number of unique haplotypes within the block (the columns in Figure 1 d). *h*_*i*_ is the dosage of the haplotype *i, h ∈* (0, 1, 2). *v* is a matrix of *c* covariates. *n* is the number of individuals. *c* is the number of covariates. In addition to haplotype-based eQTL analysis (B-Hap), eQTLHap assesses the associations between gene expression and each SNP within a block (similar to the simple linear regression model provided by Matrix eQTL), as well as the genotype of the block (B-Gen). For block’s genotype that is a categorical variable, an ANOVA regression model is fitted for this relation. The main difference between this ANOVA model and the model in Equation (1) is that here the genotype dosage is either 0 or 1. This comprehensive scan (SNP-based, B-Gen and B-Hap) was applied in all experiments reported in this study. The covariates part in Equation (1) is dropped out when such data is not included in the analysis.

SNPs with minor allele frequency (MAF) < 0.01 and haplotypes with frequency < 0.02 were eliminated to reduce the number of unique haplotypes per block. The unique haplotypes can be further reduced by considering haplotype tagging SNPs (htSNPs) as described in supplementary methods. However, tests on simulated data showed that complete haplotypes provide slightly better results as shown in supplementary figures 1 and 2, therefore, we confine the results in this manuscript to this configuration.

**Figure 2:**
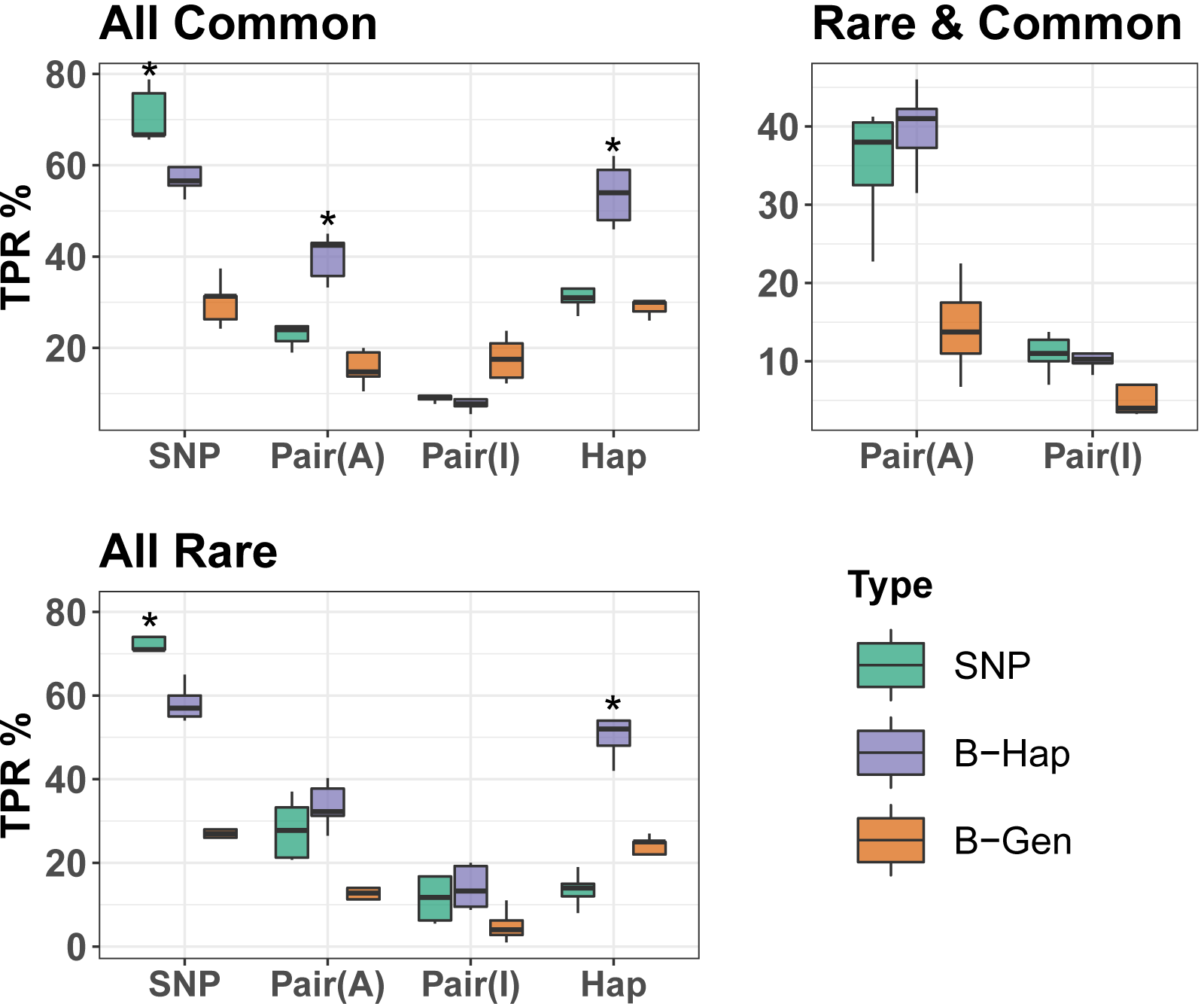
TPR for eQTL analysis based on SNP, B-Hap, B-Gen when applied to simulated genotype and gene expression data for different causal architectures. The x-axis represents the causal architecture where Pair(A) is an additive impact of a SNP pair and Pair(I) is an interaction of a SNP pair. The ‘Rare & Common’ scenario is only available for SNP pairs. Asterisks represent results that are significantly higher than all other approaches (using t-test and significance threshold of 0.05).

### 2.3 Implementation

eQTLHap is implemented in R and it depends on matrices operations to calculate correlation coefficients similar to the ultra-fast Matrix eQTL [7] to achieve high speed. It can be reconfigured to allow conducting block’s haplotype, block’s genotype and single SNP assessment individually or combined. In addition, it allows the adjustment of significance p-values through several methods accepted by *p*.*adjust* R function as well as configurable permutation analysis. Other parameters such as covariate analysis, frequency thresholds can also be adjusted for customised analysis. Further implementation details are in the supplementary methods.

### 2.4 Genotype and haplotype preparation

Haplotypes are simulated using *msprime* simulator [21] using the same configuration to that of [22]. A region of 20 mbp was simulated for 1000 individuals (regular size in eQTL studies). Diallelic SNPs and SNPs with MAF < 0.01 were then eliminated.

GEUVADIS [3] and GTEx data from https://www.ebi.ac.uk/arrayexpress/files/E-GEUV-1/ and https://www.ncbi.nlm.nih.gov/projects/gap/cgi-bin/study.cgi?study_id=phs000424.v8.p2 are used in our experiments. Quality control was applied to the genotype datasets using PLINK to keep only SNPs with MAF > 0.01, Hardy-Weinberg equilibrium (HWE) < 10^*−*6^, the missing rate per individuals < 0.1, and missing rate per SNP < 0.1. Quality controlled genotype data were phased through *consHap* consensus phase estimator [23] by aggregating 15 applications of the SHAPEIT2 tool [24]. This approach has been reported to reduce the switch error rate significantly for datasets with small sample size (< 2,000).

### 2.5 Gene expression simulation

Gene expression data were simulated for five causal genetic architectures similarly to the study [9] using the following equations:

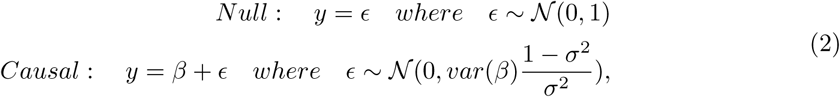

where *y* is the simulated gene expression, *β* is the assumed effect size of the underlying variant (single SNPs, pairs of SNPs or haplotypes) while *σ* is the gene expression heritability, i.e the proportion of the expression variation caused by the genetic architecture. We consider models where causal variant were common (MAF *≥* 0.05) or rare (MAF < 0.05) SNPs (*β* = *g*, where *g* is minor allele dosage of a SNP as illustrated in Figure 1 b), common and rare haplotypes (*β* = *h*, where *h* is the dosage of the haplotype) together with pairs of SNPs (additive where *β* = *g*_1_ + *g*_2_ and interaction where *β* = *g*_1_ *× g*_2_). Pairs of rare, common and a mix of rare and common SNPs were considered. In addition, we simulated expression under a null model with no variants having an effect to identify false-positive rates. *σ* was fixed at 0.05 in all simulations, with the impact of changing *σ* reported in Supplementary Figures 3-5.

**Figure 3:**
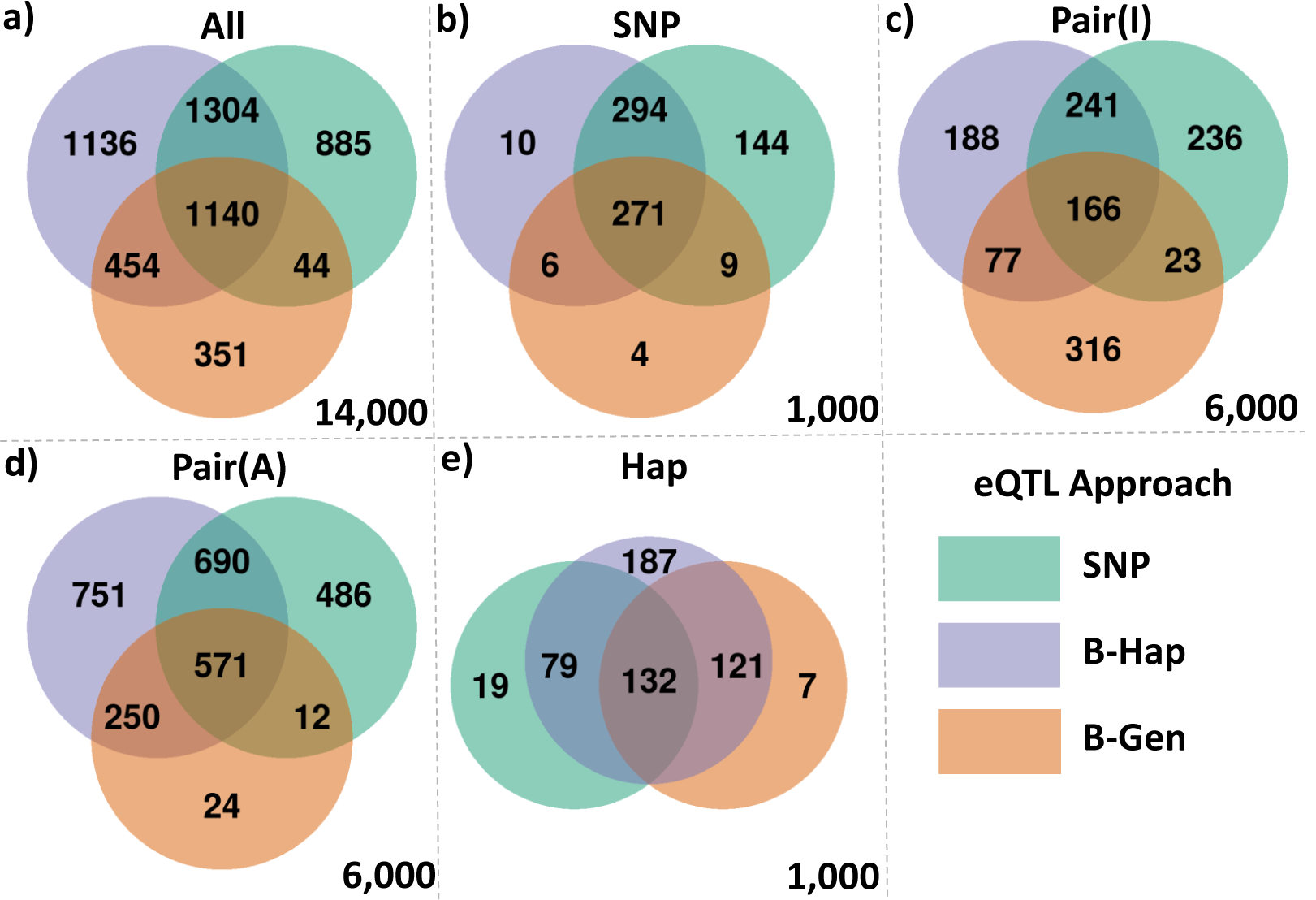
Venn diagram for significant associations detected by B-Hap, B-Gen and SNP based eQTL on simulated data. Subplots correspond to different causal architectures: a) all simulations combined. b) SNP. c) interaction of SNP pair. d) additive effect of SNP pair. e) haplotype. The total number of simulated associations are in right bottom corners.

For simulation based on a pair of SNPs, pairs within 7.5 kbp were picked (within blocks and randomly) when their correlation (squared Pearson correlation) is < 0.8 and none of them has a correlation > 0.8 with the encoding of their interaction or additive impact. Similarly, causal haplotypes were picked when they do not have a correlation > 0.8 with any single SNP within the same block. This limitation avoids the cases where these simulations will be equivalent to single SNP-based simulations. For each experiment, 100 simulations were run for each causal architecture with random initialisation. Further details of these simulations are provided within the supplementary methods.

### 2.6 Simulations to compare different eQTL approaches

To reduce the impact of synthetic data on the results, we used real genotypes for the simulations in this experiment. Genotype data were obtained from the 1000 Genome Project for 373 individuals from European population (obtained from GEUVADIS dataset). Genotypes of chromosome one were phased using SHAPEIT2 [24] and haplotype blocks were determined using PLINK software. Five genomic regions were extracted for these individuals within 1mbp up/downstream of the following arbitrarily chosen genes ENSG00000218510, ENSG00000127074, ENSG00000196539, ENSG00000162441, and ENSG00000198468. For each of the give genes, we simulate 100 common and 100 rare SNPs, leading to 1000 simulations for the single SNP and haplotype architectures. For pair based architectures, we simulate 100 pairs of common SNPs, 100 pairs of rare SNPs and 100 rare/common pairs for the give genes, and repeating simulations to consider pairings within a single block and pairings between blocks, leading to 6000 simulations in total. As such, 14,000 gene expression simulations were generated for all mentioned casual architectures. eQTL analysis is performed using *eQTLHap* for all representations (SNP, B-Gen and B-Hap).

### 2.7 Impact of phasing error on haplotype-based eQTL

To assess the impact of phasing accuracy on eQTL results, we generated five haplotype datasets from the same region with different switch errors (SE) ranging from 0 to 2.5%. We conducted eQTL analysis for each dataset using the same simulated gene expression. For this experiment, a genotype dataset was formed from simulated haplotypes by combining both haplotype copies of individuals. To obtain realistic SEs, the formed genotypes were then phased using HAPI-UR [25] 100 times and all SEs were recorded. HAPI-UR was used as it is fast and non-deterministic by default [12]. The average SE of the 100 applications was 0.78%, while the maximum SE obtained when considering all unique SE locations was 2%. Six versions of the simulated region were prepared by switching individual’s haplotypes within the locations recorded for HAPI-UR’s SEs. SE within these datasets varies from 0% (the original simulated data with no errors) to 2.5% by 0.5% step. For the version with SE = 2.5%, in addition to all recorded SE (account for 2%), we used random heterozygous SNPs as locations of SEs to reach the desired SE (2.5%).

Four different regions (1.5 mbp) were selected arbitrarily from the simulated haplotypes (SE = 0%). Haplotype blocks determined using PLINK software and overlapping with each region are used for further analysis. Gene expression data were generated as explained above. Haplotype-based eQTL analysis (B-Hap) was applied for the simulated gene expression and the 6 versions of the haplotypes (different switch errors) for each region.

### 2.8 Analysis of GEUVADIS and GTEx data

The comprehensive eQTL analysis was applied to the GEUVADIS dataset following similar configurations as its originally reported [3]. Briefly, individuals of European and Yoruba populations (373 and 89, respectively) were analysed separately. Gene expressions were used after probabilistic estimation of expression residuals (PEER) normalisation [26]. The top three principal components (PC) were used as covariates within individuals of European descent, while the top 2 PCs were used for Yoruba individuals to eliminate any impact of population stratification.

GTEx gene expression and covariates for multiple tissues were obtained from the GTEx web portal https://www.gtexportal.org/home/datasets. GTEx’s covariates datasets include hidden factors detected by PEER normalisation. For both datasets, genes with non-zero expression level for more than 90% of the individuals were investigated considering SNPs within 1mbp up/down from their transcription start site (TSS). Associations with “empirical” p-value < 0.05 were recorded for multiple test correction based on a permutation of 1,000 iterations.

### 2.9 Evaluation and comparison

After conducting eQTL analysis, p-values are reported for all associations. Multiple test correction (MTC) was carried out for all p-values (SNPs, block’s genotype, and block’s haplotypes, separately) using the Benjamini-Hochberg (BH). Associations based on simulated data are considered significant when their BH corrected p-value is < 0.05. Associations based on real data are considered significant when their BH corrected p-value is < 0.05 and the permutation-based p-value is < 0.015.

With simulated gene expression data, TPR was calculated as the percentage of detected simulated associations of all simulated associations with respect to each causal architecture. For a model to have 100% TPR for simulations based on SNP pairs, both pairing SNPs should be reported as significant. If only one SNP of each pair was detected as significant for 100 different simulations, the TPR will be 50%. For haplotype-based eQTL, the block containing the causal SNP should be reported significant. When applying SNP-based eQTL on haplotype-based simulations, if any SNP within the causal haplotype block is reported significant, the association considered detected. Venn diagrams were generated for each causal architecture, where simulations for pairs account for 2 causal SNPs.

Significant blocks reported for GTEx were transformed from GHR38 to hg19/GHR37 using the LiftOver web tool https://genome.ucsc.edu/cgi-bin/hgLiftOver to match the same genome assembly as GEUVADIS data. After that, replications between GTEx and GEUVADIS eGenes are identified when a significant block from GTEx overlaps with another one from GEUVADIS for the same gene.

## 3 Results

### 3.1 Comparison of different approaches for eQTL analysis

We compared the results of eQTL analysis using single SNPs, SNP blocks as haplotypes (denoted B-Hap) and as genotypes (denoted B-Gen) using genotypes from 273 European individuals from the 1000 Genome Project and simulated gene expression under five causal architectures. Within 5 genes, the average number of haplotype blocks is 816 with an average length of 1.25 kbp and an average SNP count of 17.

Figure 2 shows that B-Hap analysis is superior to other approaches when the causal architecture involves a haplotype stretch, an additive impact of a SNP pair or an interaction of rare SNPs. As expected, simulation-based on single SNPs are detected better via SNP-based eQTL regardless of the frequency of the SNP. B-Gen approach was more effective when dealing with interactions of common SNPs. With respect to the null causal architecture, the FDR was 0.06%, 0.05% and 0.05% for SNP, B-Hap, and B-Gen, respectively. While these results are reported for *σ* = 0.05, a similar pattern was observed when changing *σ* from 0.01 to 0.1 as shown within supplementary Figures 3, 4, and 5. The main difference is that the detection rate increased for all approaches with *σ*.

**Figure 4:**
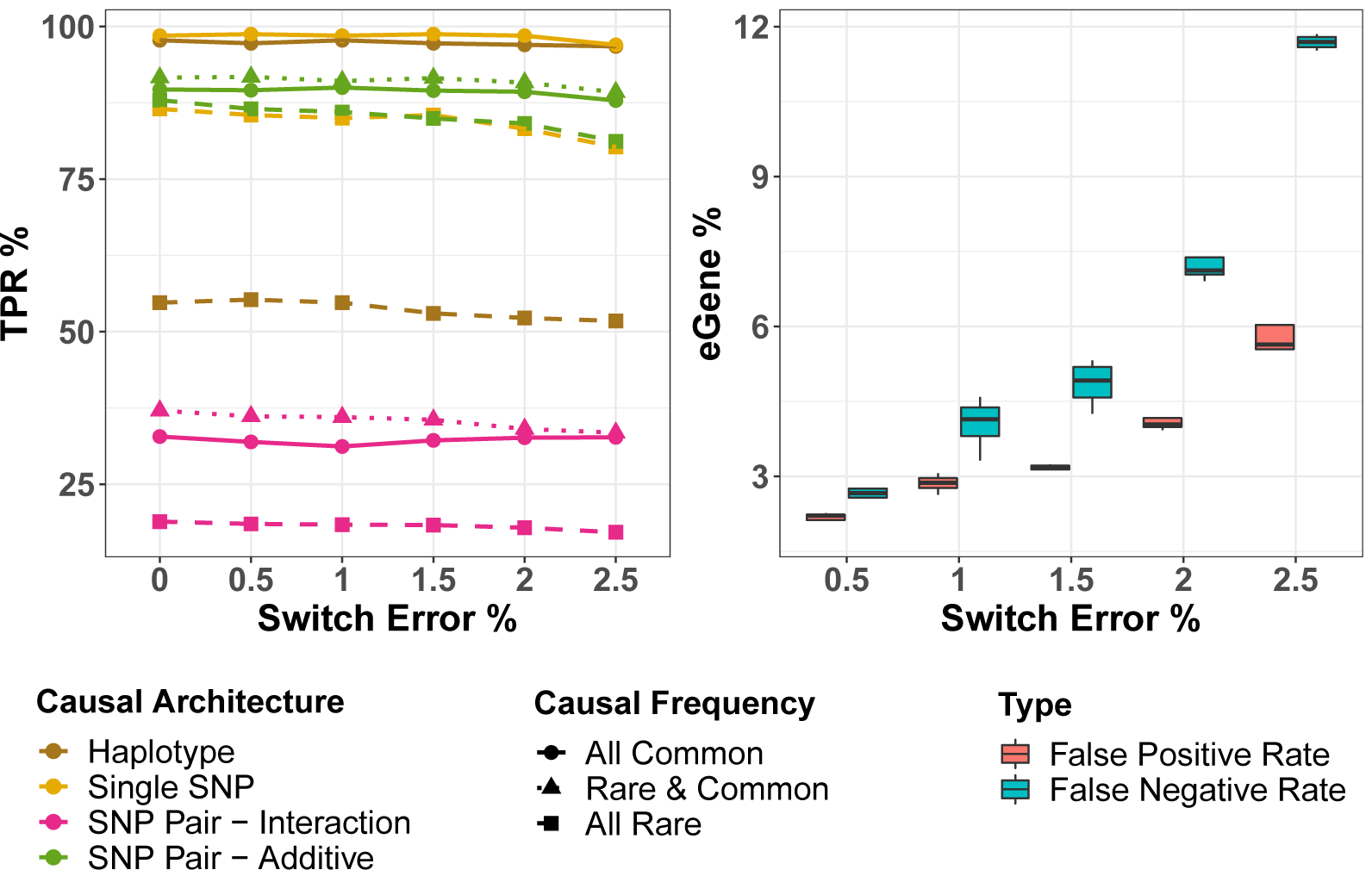
Impact of SE on haplotype-based eQTL analysis. a) TPR of the analysis with respect to switch error. b) Percentage of false positives/negatives with respect to SE.

**Figure 5:**
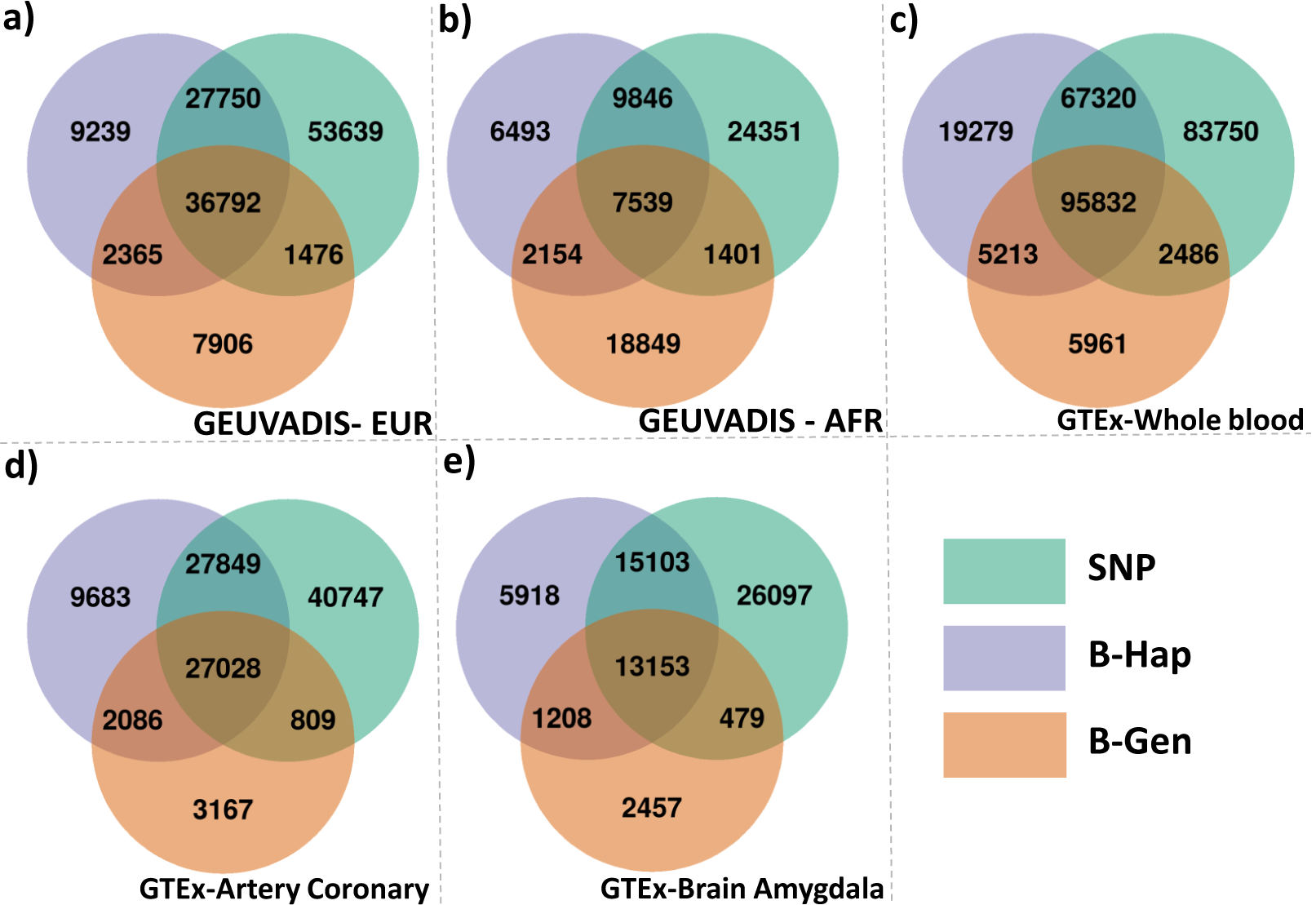
Venn diagram for detected significant associations from GTEx and GEUVADIS datasets. a) and b) are obtained from analysing 22,466 and 22,474 genes within 22 chromosomes of EUR (373 individuals) and AFR (89 individuals) populations from GEUVADIS dataset, respectively. c) is obtained from analysing 19,696 genes of GTEx-whole blood tissue (670 individuals). d) is obtained from analysing 23,735 genes of GTEx-artery coronary tissue (213 individuals). c) is obtained from analysing 23,268 genes of GTEx-brain amygdala tissue (129 individuals). These associations are considered significant as their BH-corrected p-value < 0.05 and permutation-based p-value < 0.015.

We further compared the simulated associations detected by the three approaches to quantifying the similarity and differences between these models. The Venn diagram shown in Figure 3 illustrates that each approach could identify a unique set of the simulated associations that was not captured by others, with B-Gen being the least effective method. With a SNP-based causal architecture, a SNP-based eQTL analysis has the highest power to detect associations, detecting the casual variant in 980 out of 1000 simulations. The significance of haplotype-based analysis is demonstrated with other causal architectures where this approach could reveal a large number of associations that were ignored by SNP-based analysis. We have observed from detailed results, that SNP-based analysis could detect one SNP of causal pairs for the majority of the simulations, yet, the other pairing SNPs were missing. With such cases, the haplotype-based analysis could capture both SNPs involved in the simulations. As expected, we observe a drop in TPR for all approaches when comparing results from real vs simulated genotypes.

These experiments show that the three approaches not only agree on a large percentage of the detected associations but also complement each other by revealing a unique subset of the simulated associations.

### 3.2 Impact of phasing errors on haplotype-based eQTL

In real applications, haplotype information is not perfect as they are obtained computationally through phasing methods. Therefore, it is important to assess the impact of the errors on the downstream haplotype-based eQTL analysis. Here, we assess the impact of SE, the standard metric of phasing evaluation [14], on the sensitivity or TPR of haplotype-based eQTL analysis applied to four regions of 1.5 mbp (1,945 SNPs) using simulated genotype and gene expression data.

Haplotype-based eQTL analysis was applied to an average of 108 haplotype blocks within each region. Block lengths varied from 0.03 kbp to 59 kbp with an average length equals to 12 kbp (mean SNP count is 16, min = 2 and max = 68). 6,400 simulations were generated for all causal architectures followed by association assessment for each block-gene expression pair. This analysis was repeated for 6 versions of the four regions where SE varied from 0 to 2.5% by 0.5% step. The percentage of incorrectly phased haplotype blocks within these datasets were 0.4% 0.8%, 1.1%, 1.7%, and 3.2%, respectively.

We observed that the number of reported significant associations reduced by an average of 1,495 associations when SE increased from 0 to 2.5%. After MTC, the false discovery rate (FDR) when there is no genetic causal was between 0.04% and 0.06%. Figure 4 a) shows the percentage of detected simulated associations varied from 7% to 100% depending on the causal architecture and phasing error within the data. There was a slight impact of SE within the range (0-2.5%) on the TPR of haplotype-eQTL analysis, especially when the causal architecture involves a common SNP/haplotype (solid lines in the figure). There was 3.9%, 3.6%, 2%, and 1.8% TPR reduction when SE increased from 0 to 2.5% for single SNP, an additive impact of SNP pair causal architectures, haplotype, and an interaction of a SNP pair respectively. SNP or haplotype-based association were easier to detect compared to the associations based on SNP pairs. We observed a low detection rate of simulations based on interactions of SNP pairs (less than 37%) compared to other casual architectures.

Furthermore, we investigated how the detected significant associations vary with respect to different SEs compared to the associations revealed when there were no errors present (i.e. SE = 0). Haplotype-based eQTL conducted on the data with SE = 0 reported 43,249 significant associations with a corrected p-value < 0.05. These associations represent 3% of all block-gene expression combinations tested. 11% of these associations are the true simulated associations. The comparison of these associations and the ones revealed from datasets with different SEs, shown in Figure 4 b), shows that when SE increases, the percentage of false positives (detected with SE > 0 dataset but were not detected when SE = 0) and false negatives (detected when SE = 0, but not detected when SE > 0) increases reaching 5.9% and 11.7%, respectively. However, the true simulated associations seem to be more robust against SE as the TPR illustrated in Figure 4 a) is less affected. Investigating the false positives showed that few of them were from the simulated associations (reaching 2.55%) but not captured when SE equals 0. Around 4.5% of the false negatives were from the simulated associations. 11.3% of the true positives (shared between dataset with SE = 0 and SE > 0) are true simulated associations.

These results support the application of haplotype-based eQTL analysis on real datasets where SE is usually less than 2.5% [12] especially given there is little impact on the TPR.

### 3.3 Application to GEUVADIS and GTEx data

After we demonstrated the efficacy of the approach through synthetic data, we applied the comprehensive eQTL analysis to real genotype/haplotype and gene expression dataset from both projects GEUVADIS (lymphoblastoid cell line for European and African populations) and GTEx (whole blood, artery coronary, and brain amygdala tissues).

In both GEUVADIS and GTEx, the overlap of eQTLs discovered by each representation, shown in Figure 5, shows a similar pattern to those in the simulated data. The approaches agree on a large set of associations with each approach finding a subset of unique associations. Similar trends can be seen in other tissues types (Supplementary Figure 6).

**Figure 6:**
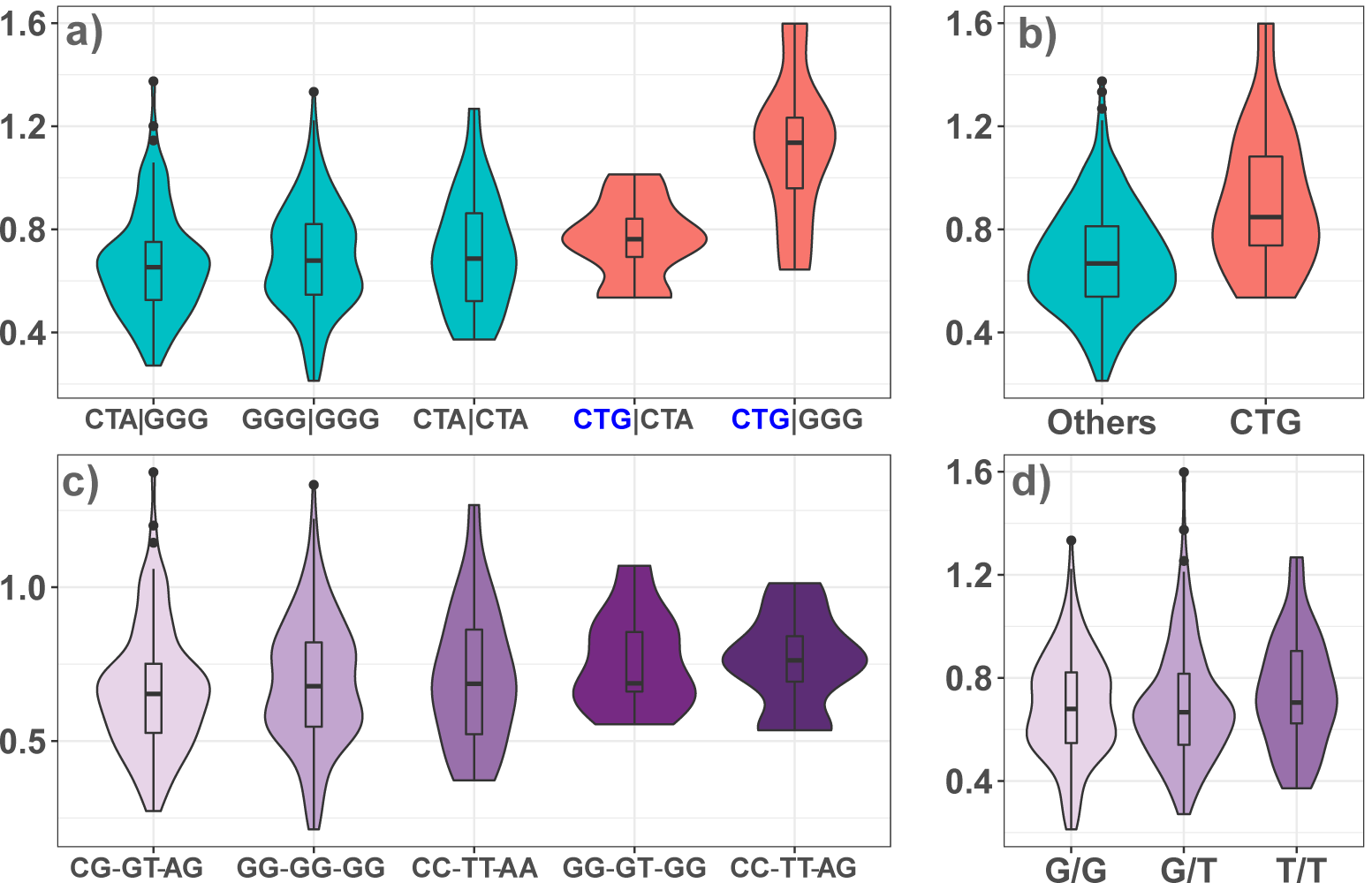
Comparison of gene expression distribution of *USP46-AS1* for a haplotype block using different representations. The block consists of three intronic SNPs (rs7657404 (MAF=0.34), rs7698053 (0.3) and rs7688816 (0.33)) on chr4:53186299-53186401. The different representations are a) phased haplotypes (q-value=0.017) b) a simplification of the strongest effect haplotype (CTG, allele frequency 0.46%) vs all others c) all genotypes (q-value =0.99) and the most significant SNP within the block (rs7657404, q-value = 0.9). p/q-values are calculated after eliminating haplotypes/genotypes with frequency *≤* 0.02 but all trends remain the same if these are included.

Applying eQTLHap to the whole blood tissue of GEUVADIS (EUR population) and GTEx, there were 14,229 (of 76,146) and 36,696 (of 187,644) significant haplotype-based associations whose p-values are less than 100 times the p-value of both block’s genotype and each SNP within the same block. These results include 1,035 and 748 genes that only haplotype-based eQTL could detect significant associations with. The average p-value of these eGenes is 2*×*10^*−*4^ and 2*×*10^*−*4^ within GTEx and GEUVADIS, respectively.

Figure 6 shows an example of a haplotype-only association, plotting the distribution of expression for *USP46-AS1* in whole blood from the European population of GEUVADIS across different variant representations for a block of three intronic SNPs (rs7657404, rs7698053, rs7688816). In this instance, the haplotype analysis is highly significant (q-value< 0.017), while the single SNP and genotype block representations are not (q-values 0.9 and 0.99, respectively). Simplifying the haplotype analysis in Figure 6 b), we see that the individuals carrying the CTG haplotype have substantially higher gene expression than those who do not. These three SNPs were found in GTEx to be significant eQTLs for RASL11B in Testis and DANCR in cultured fibroblasts [27] but we believe this to be the first association with *USP46-AS1* in whole blood reported to date.

Furthermore, we searched for replicable, significant associations from our haplotype-based eQTL analysis from GTEx and GEUVADIS dataset for whole blood tissue. There were 50,616 common associations for 2,425 unique genes reported for both datasets with 3,136 common genes between both datasets. The average p-value of these common associations is 4×10^*−*4^ and 9×10^*−*4^ with respect to both GTEx and GEUVADIS, respectively. From these 2,425 genes, there were 11 common eQTLs for 7 eGenes (Supplementary Table X) that only had significant associations when they were represented using B-Hap. To further validate these findings, we compared these associations with eQTL results those of previous meta-analyses [28, 29] finding our 11 eQTLs overlap with significant findings reported in at least one of these substantially larger studies, highlighting the increased power of our proposed approach for certain eQTLs.

We also searched for eQTLs that were detectable using haplotype but not single SNPs that could be replicated across tissues in the GTEx dataset. In total (including haplotypes that are detected using single SNPs), we find 13,573 common associations (1,312 unique genes) across whole blood (187,644 associations), artery coronary (66,646 associations), and brain amygdala (35,382 associations) tissues. The averaged p-value for these associations respectively to the mentioned tissue order is 1×10^*−*4^, 4×10^*−*4^, and 7×10^*−*4^ indicating that many of these are highly significant. Of these, 729, 1085, and 947 eGenes were only detected by haplotype-based eQTL applied to whole blood, artery coronary, and brain amygdala, respectively. 11 eGenes of them replicate in across at least one other tissue type.

These findings demonstrate that our haplotype approach is able to uncover novel eQTLs that are undetectable by single SNP approaches, with a subset replicating across either dataset or tissue. These results further highlight the utility of phase-aware haplotypes for eQTL analysis.

## 4 Conclusion

In this study, we propose *eQTLHap*, a haplotype-based eQTL approach at a block scale that serves as a complementary analysis to genotype-based eQTL.

Our results show that haplotype-based eQTL outperformed other approaches for eQTL when the causal genetic architecture comprises multiple SNPs. According to the results obtained in this study, the three approaches of eQTL (based on SNP, block’s genotype and block’s haplotype) agreed on a large proportion of the detected associations. At the same time, each approach captured a unique subset of the associations that have not been detected by other approaches. This observation shows that the different approaches complement each other and can provide a comprehensive eQTL scan when applied together.

To the best of our knowledge, this is the first study that investigates how phasing errors affect the results of downstream eQTL analysis. Experiments applied to synthetic haplotype and gene expression datasets demonstrated the small impact of phasing errors on TPR of downstream eQTL findings (SE ¡ 2.5%). However, there was variation in all reported significant associations demonstrate by increased false positive and negative rates when SE increased from 0% to 2.5%. This impact can be mitigated by improving phasing accuracy using consensus estimators [12] or phasing the data multiple times, applying eQTL analysis to each and keeping the stable findings.

The higher TPR obtained by haplotype-based eQTL can be justified by the less conservative MTC applied to block assessment compared to single SNPs as block count is substantially less than SNP count. However, the fact that haplotype-based eQTL outperformed block’s genotype eQTL analysis in most of the experiments demonstrates that this enhanced performance is also associated with including haplotype information in the analysis, as both assessments are applied to the same blocks.

The findings when applying eQTLHap to real dataset demonstrate the efficacy of this approach as there was a large agreement with standard SNP-based eQTL approaches. In addition, several results were only revealed when considering haplotype information which replicated in both GTEx and GEUVADIS datasets as well as independent analyses available through previous meta-analyses. An interesting avenue of future research would be to explore the properties of these haplotype-based eQTLs to understand whether they represent examples of phase-specific regulation of expression or whether the findings are due to changes in the statistical test. In either case, the findings in this study highlight the potential of haplotype eQTL analysis to uncover eQTLs that have been missed through standard SNP based analysis.

eQTLHap provided in this study is simplified to accept several configurations in order to control and customise the analysis based on the user’s preference. eQTLHap allows applying SNP-based, block’s genotype/haplotype-based analysis separately and combined (with/without covariates). It accepts any kind of haplotype blocks. MTC can be applied using several standard methods, as well as permutation-based correction. Finally, all thresholds and options used for filtration and other purposes can be tuned.

## Acknowledgements

The authors gratefully acknowledge the GEUVADIS study. The Genotype-Tissue Expression (GTEx) Project was supported by the Common Fund of the Office of the Director of the National Institutes of Health, and by NCI, NHGRI, NHLBI, NIDA, NIMH, and NINDS. The data used for the analyses described in this manuscript were obtained from dbGaP accession number phs000424.v8.p2 on 12/13/2018.

