## Supplementary Materials for "A novel haplotype-based eQTL approach identifies genetic associations not detected through conventional SNP-based methods"

Ziad Al Bkhetan, Gursharan Chana,
Cheng Soon Ong, Benjamin Goudey, Kotagiri Ramamohanarao

### 1 Supplementary methods

#### 1.1 Haplotype block determination and representation

Haplotype blocks are determined based on linkage disequilibrium (LD). PLINK software [1] that implements Confidence Interval "CI" algorithm [2] is utilised for this purposes using its default parameters that limit the calculation to SNP pairs within 20kbp distance. A pair is considered in a strong LD if the bottom of the 90% D-prime confidence interval is greater than 0.70, and the top of the confidence interval is at least 0.98. These configurations can be changed using PLINK options listed at this link `https://www.cog-genomics.org/plink/1.9/ld#blocks`. All SNPs between the first and last SNPs of each block are considered as long as their MAF is greater than 0.01. We used this approach as it is reported to maintain high accuracy of the phased haplotypes at block scale (demonstrated in the third chapter).

Previous studies used specific SNPs to represent the haplotypes for eQTL and GWAS studies termed haplotype tagging SNPs (htSNPs) [3, 4]. In this study, different haplotype templates based on different htSNPs were determined within each block through an iterative procedure described below:

1. SNPs with $MAF = 0$ are removed as they do not add to the associations.
2. Pearson correlation is calculated for each pair of SNPs within the block using phased alleles.
3. For each pair of SNPs with correlation equal to 1, an arbitrary SNP is eliminated as such SNPs do not affect haplotype diversity within the block.
4. In an iterative procedure that stops when the unique haplotypes based on the chosen htSNPs are $\geq \alpha$ of the unique haplotypes based on all SNPs, do the following:
   (a) Pick the pair of SNPs with the highest correlation.
   (b) Remove one SNP of this pair that has the highest correlation with the remaining SNPs. As we want to keep the most informative SNPs that differentiate between haplotypes.

Four different thresholds were used on simulated data: 70% and 80%, 90% and 100% (complete haplotype).

#### 1.2 Gene expression simulation

Gene expression data were simulated for comprehensive evaluation and comparison between different approaches for eQTL analysis similar to the studies [5]. 11 different scenarios were used for the simulation using the following general equation:

$$y = \beta + \epsilon \quad where \quad \epsilon \sim \mathcal{N}(0, var(\beta)\frac{1-\sigma^2}{\sigma^2}) \tag{1}$$

$\sigma^2$ represent the proportion of the variance in the gene expression that can be explained by a specific causal genetic architecture[5]. In this study, 10 different values were investigated (from 1% to 10%, by 1% step). Results for $\sigma = 5\%$ were reported in the manuscript. $\beta$ represents the causal architecture encoding that can be calculated for each simulation type as follow:

1. **Null:** This simulation represent the cases where there is not genetic architecture for influencing the gene expression.
$$\beta = 0 \quad and \quad \epsilon \sim \mathcal{N}(0, 1) \tag{2}$$
With respect two this causal architecture we simulated two scenarios: Common causal SNPs when the SNPs are chosen when their MAF $\geq$ 0.1 and for rare causal SNPs with 0.02 $\leq$ MAF $\leq$ 0.05. common and rare SNPs used in all simulations are determined using the same thresholds mentioned here.
2. **Single causal SNP:** the encoding of the causal SNP used for the simulation is the scaled minor allele dosage of the SNP (0, 1, and 2).
$$\beta = scale(g_i) \quad ; \quad g_i \in [0, 1, 2] \tag{3}$$
With respect two this causal architecture we simulated two scenarios: Common causal SNPs when the SNPs are chosen when their MAF $\geq$ 0.1 and for rare causal SNPs with 0.02 $\leq$ MAF $\leq$ 0.05. common and rare SNPs used in all simulations are determined using the same thresholds mentioned here.

3. **Additive impact of two SNPs:** The encoding of causal architecture is calculated based on the genotypes of a random pair as follow:

$$\beta = scale(g_i) + scale(g_k) \quad ; \quad g_i \quad and \quad g_k \in [0, 1, 2] \tag{4}$$

   Three scenarios were simulated with respect to this architecture: two common SNPs, two rare SNPs, and a pair of common and rare SNPs.

4. **SNPs interactions:** The encoding of causal architecture is calculated based on the genotypes of a random pair as follow:

$$\beta = scale(g_i) \times scale(g_k) \quad ; \quad g_i \quad and \quad g_k \in [0, 1, 2] \tag{5}$$

   Three scenarios were simulated with respect to this architecture: two common SNPs, two rare SNPs, and a pair of common and rare SNPs.

5. **Causal haplotype:** the encoding of the causal haplotype is calculated similarly to the single SNP architecture. Instead of the minor allele dosage, we used the dosage of a specific haplotype within the paternal and maternal copies of each block. Blocks are determined using plink software as mentioned in 1.1 Haplotype block determination and representation. Haplotypes are chosen not to be highly correlated with any single SNP within the same block ($r < 0.8$).

$$\beta = scale(h_i) \quad ; \quad h_i \in [0, 1, 2] \tag{6}$$

   Two scenarios were simulated for this architecture: rare and common haplotypes using the same thresholds for rare and common SNPs.

When the causal architecture of any simulation involves two SNPs ($S1$, $S2$), either additive impact or interaction, the following conditions are satisfied:

1. Genomic distance between the SNPs is $\leq 7.5$ kbp.
2. Pairs are identified to be in the same blocks in 50% of the simulations, and also to be picked randomly in the other 50%.
3. Squared Pearson correlation between $S1$ and $S2$ is $< 0.8$.
4. With respect the encoding of the pair $\beta$, Pearson correlation between $S1$ and $\beta$ as well as $S2$ and $\beta$ is $< 0.8$.

Pair simulations were also repeated with the same conditions above, but forcing the pairs to be within the same haplotype block. Haplotypes for simulations were also picked when the Squared Pearson correlation between the causal haplotype and all SNPs within the same block is $< 0.8$. These constraints avoid the scenarios where these simulations are equivalent to single SNP simulations.

#### 1.3 Comprehensive eQTL analysis

eQTL analysis was conducted in this study for each gene-block pairs with respect to the block's genotype encoding, haplotype encoding and each SNP separately as follow:

1. **Gene expression-SNP association** The association was assessed between the gene expression and each SNP with a block separately. Each SNP is assumed to have an additive impact encoded based on the minor allele dosage (0, 1, and 2) (illustrated in Figure 1 in the manuscript - b)). Only SNPs with MAF $> 0.01$ are considered for this assessment. A linear regression model (Equation 7) is used to represent the relation. With respect to the example mentioned in Figure 1 in the manuscript - b) there will be four different models as follows: $y \sim s_1$, $y \sim s_2$, $y \sim s_3$, $and\, y \sim s_4$. The significance of each relation is calculated using t statistics and p-value.

$$y = \beta s + \alpha \tag{7}$$

   Where $s$ is the dosage of the minor allele within the SNP , $s \in (0, 1, 2)$ .

2. **Gene expression-block's genotype association** The genotypes of SNPs are combined together into one value representing the genotype of the whole block as illustrated in Figure 1 in the manuscript - c). Analysis of variance (ANOVA) model is used to represent the relation between gene expression and genotype-based

encoding of a block. F-test and p-values are calculated based on the ANOVA model and reported for the association. The ANOVA model is equivalent to multiple linear regression model:

$$y = \sum_{i=1}^{m} \beta_i g_i + \alpha \tag{8}$$

Where $m$ is the number of unique genotypes within the block. $g_i \in (0, 1)$ represents whether the individual carries the genotype $g_i$ or not. Rare genotypes within each block were eliminated from the model (genotype frequency $\leq 0.02$). Considering the same example in Figure 1 in the manuscript - c), the model is

$$y = \beta_1 g_1 + \beta_2 g_2 + \beta_3 g_3 + \beta_4 g_4 + \alpha \tag{9}$$

For a block's genotype $g$, $g_1 = I(g = 1210)$, $g_2 = I(g = 1220)$, $g_3 = I(g = 1211)$ and $g_4 = I(g = 2200)$.

3. **Gene expression-block's haplotype association** Haplotypes are encoding similarly to bag-of-words representation using in text mining. unique haplotypes within a block and across all individuals are determined then individual's block is represented using the dosage of each haplotype within both homologous chromosome copies as illustrated in Figure 1 in the manuscript - d). A multiple linear regression model is used to assess the gene expression as a response to the additive model of the haplotypes within each block.

$$y = \sum_{i=1}^{m} \beta_i h_i + \alpha \tag{10}$$

Where $m$ is the number of unique haplotypes within the block. $h_i$ is the dosage of the haplotype $i$, where $h \in (0, 1, 2)$. F statistic and p-value is calculated for the whole model. Rare haplotpes within each block were eliminated from the model (haplotype frequency $\leq 0.02$). Considering the example in Figure 1 in the manuscript - d), the model is

$$y = \beta_1 h_1 + \beta_2 h_2 + \beta_3 h_3 + \alpha \tag{11}$$

For block's hapotypes $\bar{h_1}$ and $\bar{h_2}$:
$h_1 = I(\bar{h_1} = 0110) + I(\bar{h_2} = 0110)$,
$h_2 = I(\bar{h_1} = 1100) + I(\bar{h_2} = 1100)$, and
$h_3 = I(\bar{h_1} = 0111) + I(\bar{h_2} = 0111)$.

Before conducting any statistical assessment, SNPs dosage, haplotype dosage, and gene expression were standardised as used in a well-known eQTL analysis tool (Matrix eQTL [6]).

#### 1.4 Consideration of covariates

It is popular to account for covariates in eQTL analysis such as individual's gender, imputed SNPs, principle components determined for individuals genotypes (to avoid population stratification). For such cases, the regression models above become:

$$y = \beta s + \sum_{j=1}^{c} \gamma_j v_j + \alpha \tag{12}$$

$$y = \sum_{i=1}^{m} \beta_i g_i + \sum_{j=1}^{c} \gamma_j v_j + \alpha \tag{13}$$

$$y = \sum_{i=1}^{m} \beta_i h_i + \sum_{j=1}^{c} \gamma_j v_j + \alpha \tag{14}$$

where $c$ is the covariates number. The corrected degrees of freedom for the model decreases by $c$ to account for the added covariates.

#### 1.5 Technical implementation of the statistical assessment

Statistical assessment for eQTL analysis is computationally expensive due to a large number of possible gene expression-genomic locus pairs. For fast execution, we adopted Matrix eQTL approach [6] that relies on matrix operations.

For a haplotype block for $n$ individuals and covariate matrix ($v$) of $c$ covariates, the applied algorithm (adapted from Matrix eQTL to suit blocks instead of SNPs) is described below:

1. Create bag of haplotype (*boh*) data structure for the block as a list of $m$ one-row matrices of a length $n$.
2. Fill each matrix with the dosage of its associated haplotype within the paternal and maternal copies.
3. Center all matrices: gene expression ($y$), each matrix of the bag of haplotype in (1), and the covariates ($v$) to remove the intercept ($\alpha$) from the model.
4. Orthogonalise the gene expression with respect to the covariates.

$$\tilde{y} = y - \langle y,\, v \rangle v \tag{15}$$

Where $\langle y,\, cov \rangle$ is the cross product of $y$ and $cov$.

5. Orthogonalise each matrix ($h_i$) of *hob* with respect to the covariates and other matrices.

$$\ddot{h_i} = h_i - \langle h_i,\, v \rangle v \quad ;\, i = 1, 2, \ldots, m. \tag{16}$$

$$\tilde{h_i} = \ddot{h_i} - \langle \ddot{h_i},\, \tilde{h_j} \rangle h_j \quad ;\, i = 1, 2, \ldots, m; \quad j = 1, \ldots, i-1. \tag{17}$$

6. Standardise each matrix ($\tilde{h_i}$) of *hob*.
7. Calculate test statistic $R^2$:

$$R^2 = \sum_{i=1}^{m} \langle \tilde{y},\, \tilde{h_i} \rangle^2 \tag{18}$$

8. Calculate F-test then pvalue based on $R^2$

$$\text{F-test} = \frac{(n - m - c)R^2}{m(1 - R^2)} \tag{19}$$

In case no covariates provided, the same algorithm is applied by ignoring the steps that involve the covariates. For SNP-based eQTL, the same algorithm as Matrix eQTL is applied considering the simple linear regression model.

The same algorithm can be applied to assess the association between gene expression and the genotype of the block. The only difference is to change boh matrices to include the dosage of the genotypes.

### 2 Supplementary results

#### 2.1 Different templates for haplotype blocks

We investigated different haplotype templates for association assessment to find the template that leads to the best results. For this purpose, htSNPs were chosen in a way they represent 70 %, 80% and 90% of all haplotypes determined using all SNPs within each block. Using simulated data with different switch error rates, we found that different templates obtained similar TPR as illustrated in figure 1. For further investigation, we compared

Figure 1 **TPR of haplotype-based eQTL using different haplotype templates**. TPR calculated haplotype templates representing 100%, 90%, 80% and 70% of blocks haplotypes with respect to different causal architecture and causal frequency. Each plot represents a TPR comparison for different switch error rate from 0 to 2.5%. Results are summarised for all causal architectures mentioned in the section 2.5 Gene expression simulation in the manuscript.

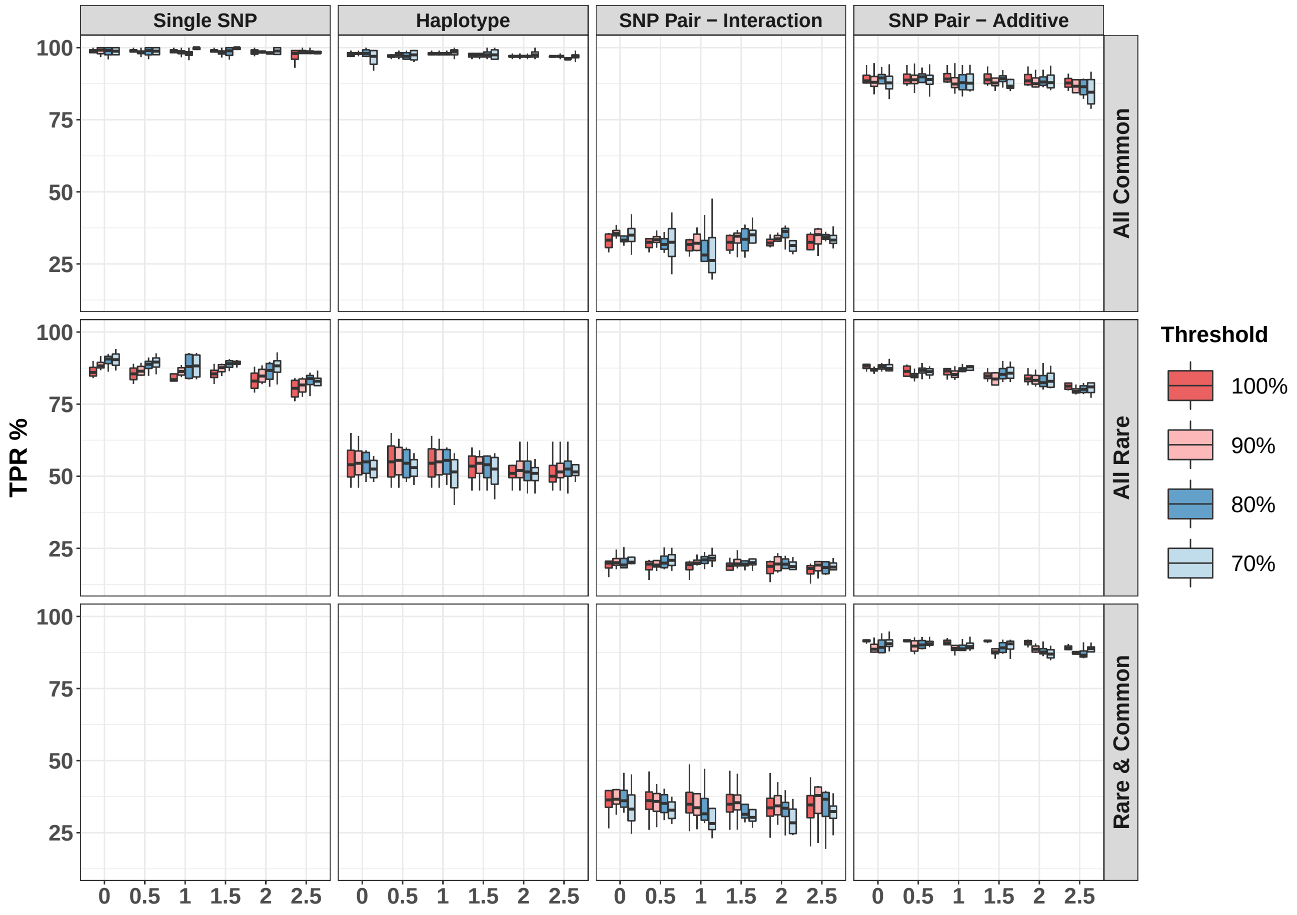

results' variation for different SEs and with respect to each template. Compared to the associations detected on haplotypes without errors (SE = 0%), using all SNPs within a block maintained more similar results than other templates as illustrated in Figure 2. Both missing eGenes (only reported on data with SE = 0%) and False eGenes (only reported on data with errors) increase when the SNPs within each block decrease. Moreover, The difference between these templates increases with the switch error rate. These observations encouraged to use 100% template for the majority of the results in this chapter.

Figure 2 **False positive and false negative rate for different haplotype templates.** A comparison of the percentage of false positive/negative rate for different haplotype templates and switch errors. These results represent a comparison to the eGenes obtained on the data without errors (SE = 0%). Results are summarised for all causal architectures mentioned in the manuscript.

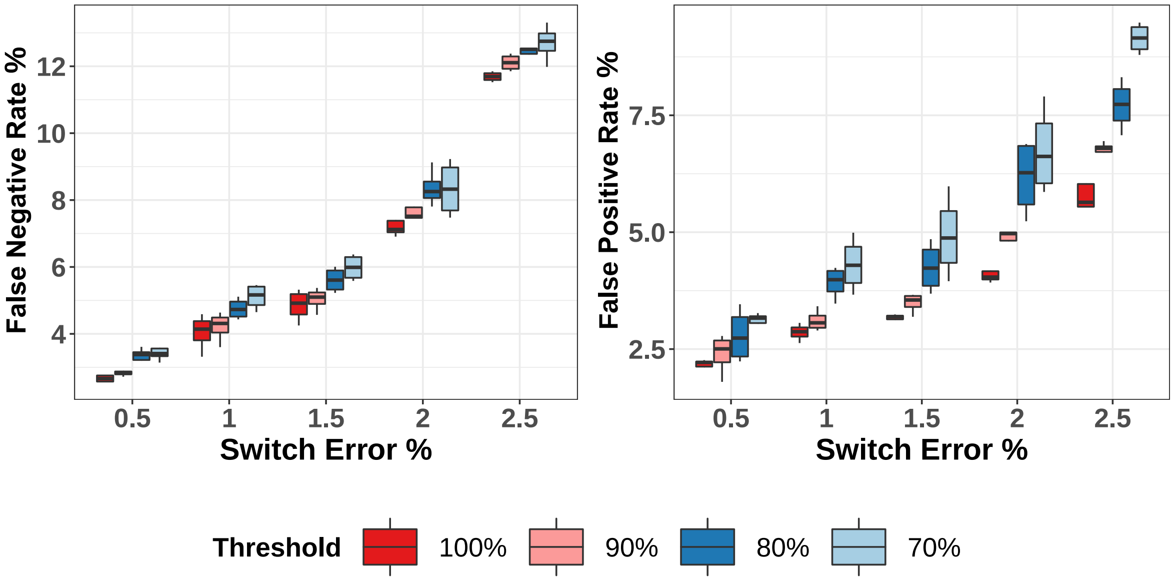

#### 2.2 Performance of eQTL approaches with different gene expression heritablity

In this section, we report the performance of eQTL approaches with respect to different values of $\sigma$ from 0 to 0.1 where $\sigma$ is the proportion of the variance in the gene expression that can be explained by a specific causal genetic architecture. This experiment was applied to genotype data obtained from the 1000 genome project for 373 individuals from European population and gene expression data simulated for the genes ENSG00000127074, ENSG00000162441, ENSG00000196539, ENSG00000218510, ENSG00000198468 genes as used in section 3.1 Comparison of different approaches for eQTL analysis in the manuscript. As expected the TPR increases with $\sigma$ as illustrated in Figures 3, 4 and 5 that represent causal architectures with different frequencies.

eQTL based on block's genotype obtained the minimal TPR for all experiments with an exception the one simulated based on haplotypes. SNP-based eQTL outperformed all other approaches when the causal architecture is a single SNP regardless of the $\sigma$. Haplotype-based eQTL analysis is superior to other approaches when the causal architecture involves multiple SNPs. When the causal is a pair of common SNPs, the improvement in TPR obtained by haplotype-based eQTL increases with $\sigma$ as illustrated in Figures 3, 4 and 5. TPR patterns for all approaches are similar regardless of the frequency of causal architecture.

Figure 3 TPR of eQTL approaches for different common causal architectures with respect to different values of gene expression heritability.

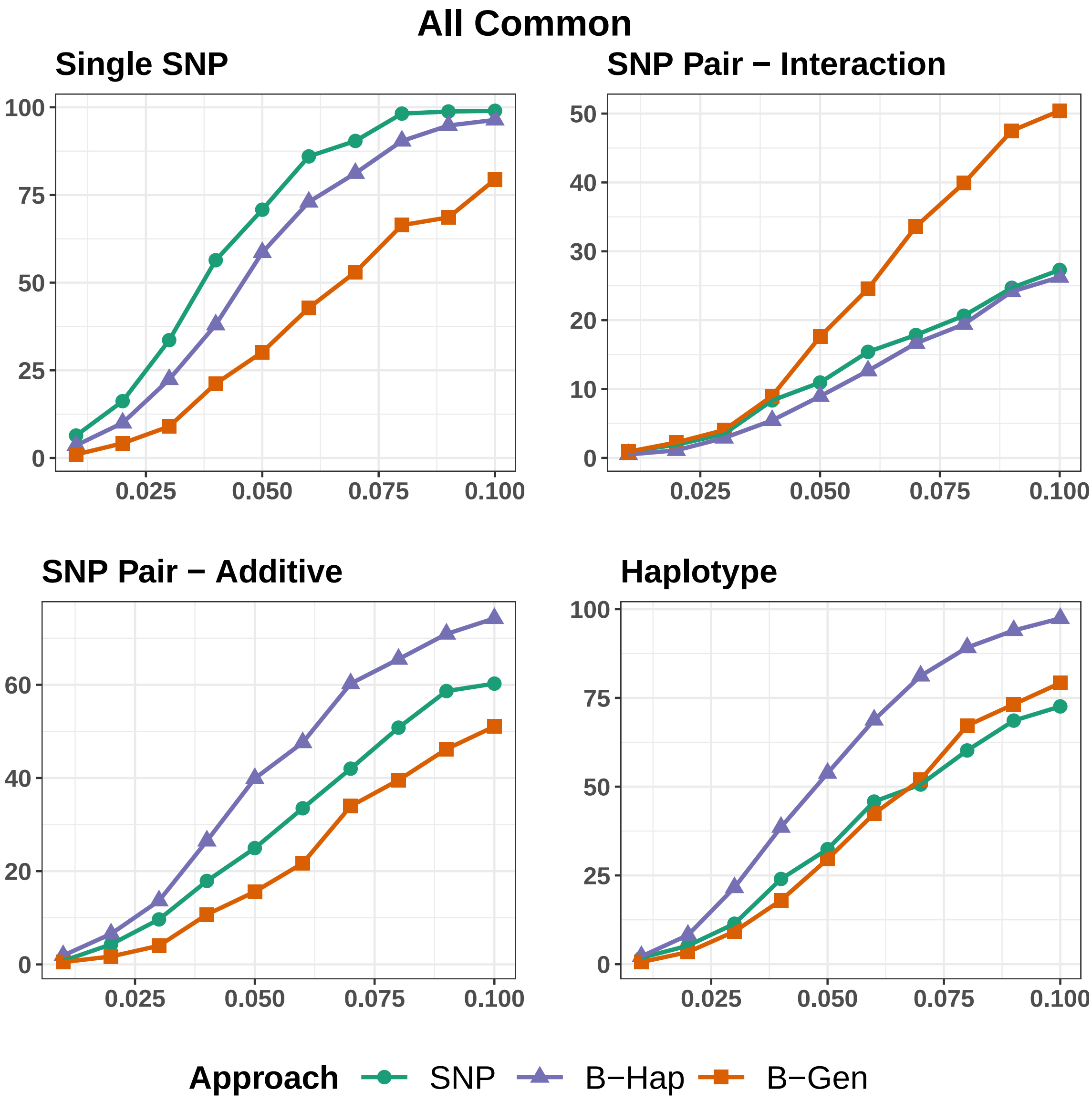

Figure 4 TPR of eQTL approaches for different rare causal architectures with respect to different values of gene expression heritability.

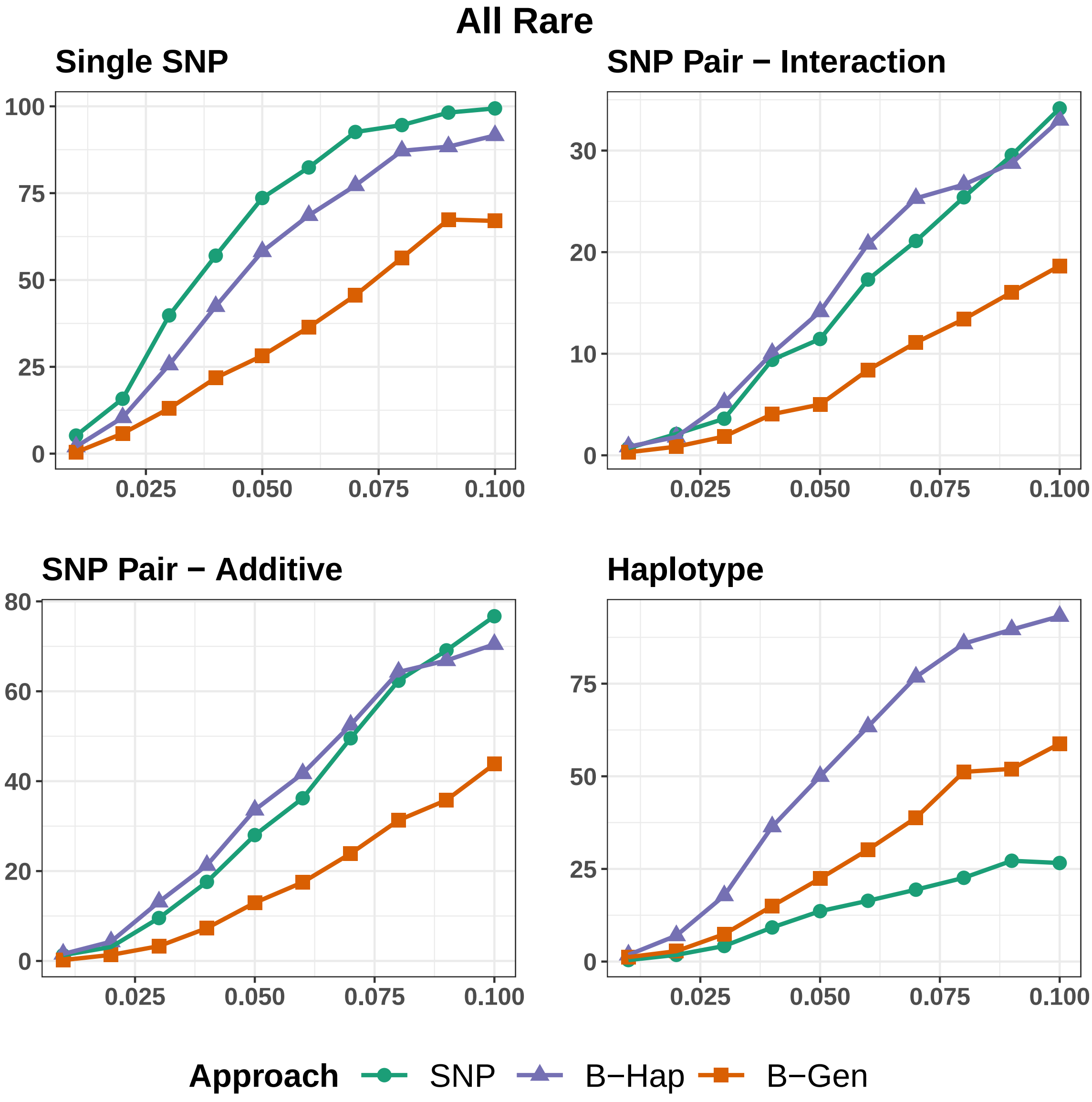

Figure 5 TPR of eQTL approaches for causal architectures based on a pair of rare and common SNPs with respect to different values of gene expression heritability.

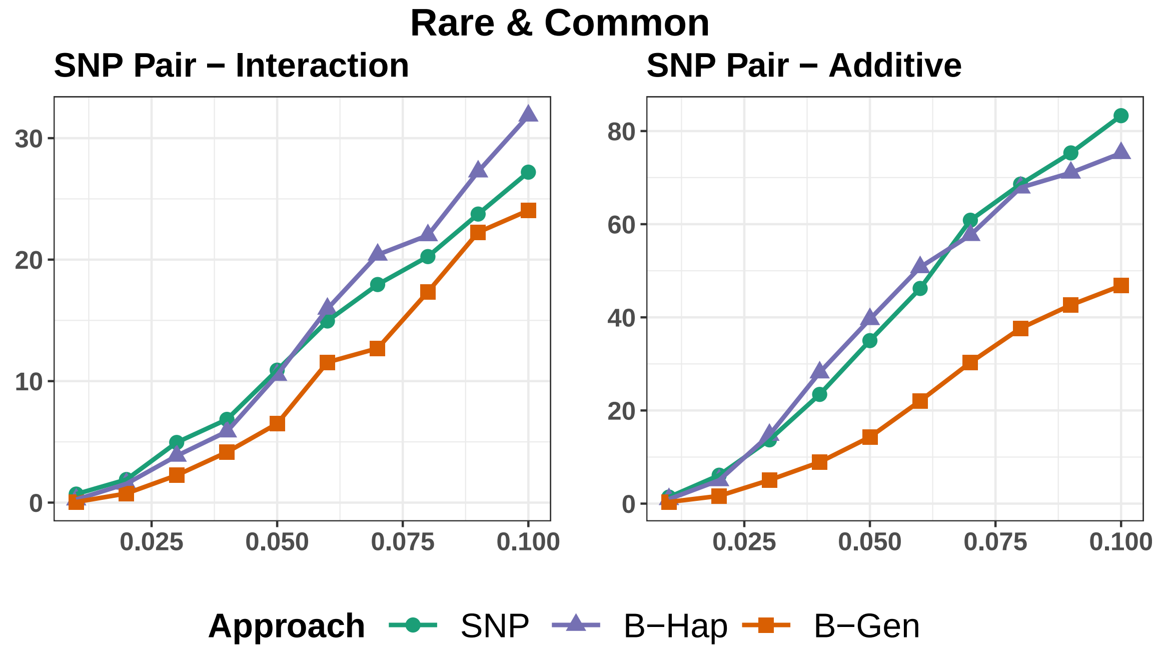

Figure 6 Venn diagram for detected significant associations from different tissues provided by GTEx datasets. a) is obtained from analysing 24,724 genes of GTEx-adipose visceral omentum tissue (469 individuals). b) is obtained from analysing 25,849 genes of GTEx-breast mammary tissue (396 individuals). c) is obtained from analysing 22,759 genes of GTEx-cells EBV-transformed lymphocytes tissue (147 individuals). d) is obtained from analysing 25,479 genes of GTEx-spleen tissue (227 individuals). e) is obtained from analysing 24,290 genes of GTEx-stomach tissue (324 individuals). f) is obtained from analysing 25,873 genes of GTEx-nerve tibial tissue (532 individuals).

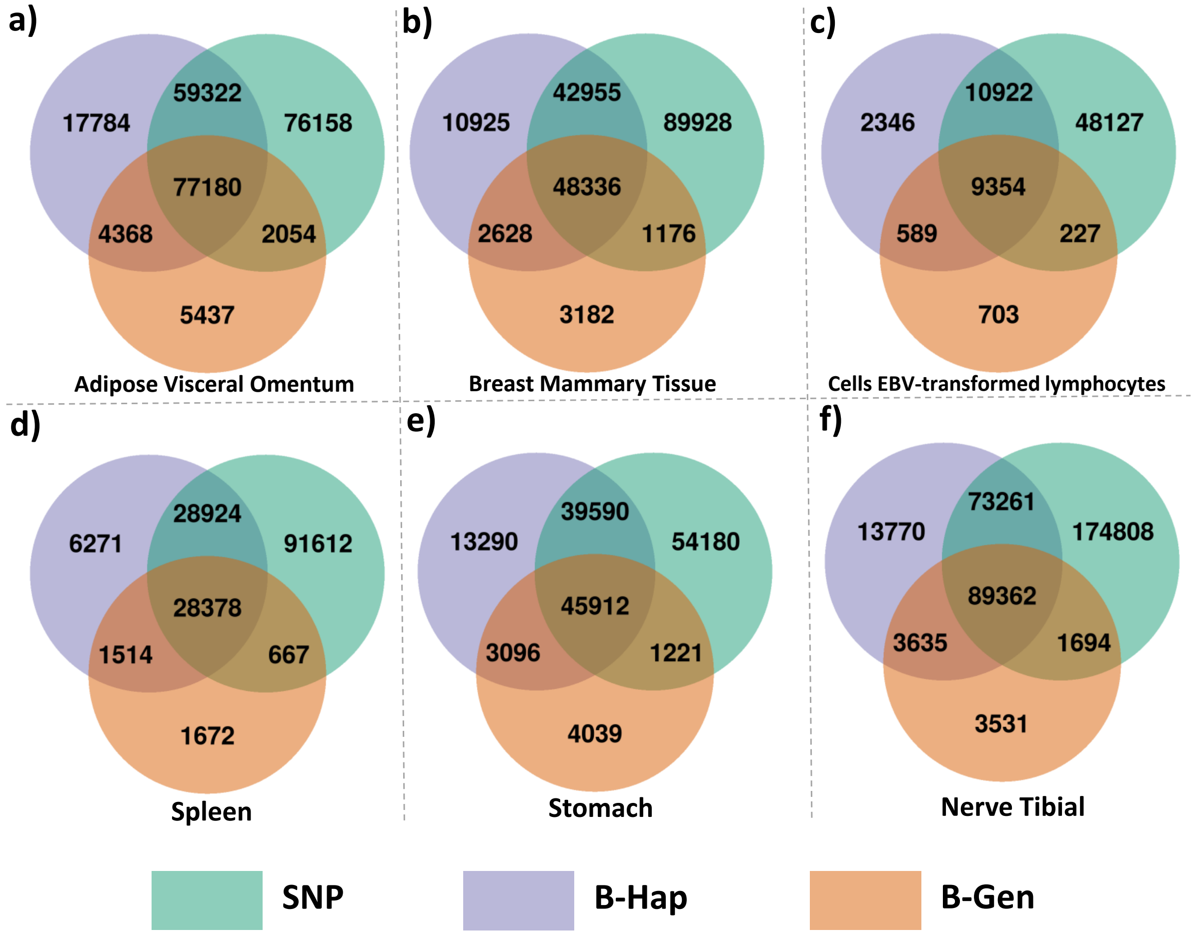
